# DataJoint: managing big scientific data using MATLAB or Python

**DOI:** 10.1101/031658

**Authors:** Dimitri Yatsenko, Jacob Reimer, Alexander S. Ecker, Edgar Y. Walker, Fabian Sinz, Philipp Berens, Andreas Hoenselaar, R. James Cotton, Athanassios S. Siapas, Andreas S. Tolias

## Abstract

The rise of big data in modern research poses serious challenges for data management: Large and intricate datasets from diverse instrumentation must be precisely aligned, annotated, and processed in a variety of ways to extract new insights. While high levels of data integrity are expected, research teams have diverse backgrounds, are geographically dispersed, and rarely possess a primary interest in data science. Here we describe DataJoint, an open-source toolbox designed for manipulating and processing scientific data under the relational data model. Designed for scientists who need a flexible and expressive database language with few basic concepts and operations, DataJoint facilitates multiuser access, efficient queries, and distributed computing. With implementations in both MATLAB and Python, DataJoint is not limited to particular file formats, acquisition systems, or data modalities and can be quickly adapted to new experimental designs. DataJoint and related resources are available at http://datajoint.github.com.

## Introduction

Data emerging from today's biological experiments are not merely “big” but increasingly multimodal and dynamic, as projects quickly move to new technologies and experimental paradigms [1–6]. In our field of neuroscience, a single experiment may involve several signal acquisition modalities such as imaging, multi-channel electrophysiology, genetic sequencing, optical stimulation, behavioral recordings, and an array of other techniques [7,8].

Concerted effort must be applied to maintain the integrity and reproducibility of scientific findings by keeping data intelligible and uncorrupted over months and years of experiments. Data shared between research groups are particularly susceptible to confusion and corruption during data transfers and concurrent access. Data integrity must be preserved when it is accessed by multiple computers performing parallel and distributed computations in scenarios such as cluster or cloud computing, even when some jobs are interrupted.

Flexible data access can greatly increase productivity during data analysis to obtain specific cross-sections stored data based on diverse criteria from multiple datasets as in the case of summary statistics across multiple experiments. When using the file system for organizing data, such analysis may require traversing folders and files, parse their contents, and select and assemble the necessary data for each analysis [4]. Changing experimental configurations require careful adaptations in the structure of associated data repositories and tedious reconfiguration of analysis scripts.

In contrast to file systems, relational databases explicitly maintain data integrity and offer flexible access to crosssections of the data [9]. A relational database preserves consistency and referential integrity even as multiple users and processes manipulate data concurrently or if a process terminates unexpectedly. Unlike a file system, a database is not an inert repository designed to simply reproduce data in their original form: instead, it provides a way to access specific and precise cross-sections of the data in order to answer questions unforeseen at the time the data are deposited. A database system also defines and enforces real-world constraints and relationships between data elements, even as experimental paradigms change over time. It communicates and enforces these relationships because they are inherent in the structure of the data itself. This form of self-documentation enables new users to readily understand the architecture of an unfamiliar data set.

Here we describe an open-source framework, DataJoint, to help scientists organize, populate, and query large volumes of data. In contrast to existing database systems and query languages which are overgrown with extraneous complex [10, 11], DataJoint introduces a minimal set of operators that allow flexible, efficient, and expressive queries to retrieve exactly the data one needs. Unlike other domain-specific data management tools [11–17], DataJoint is general and extensible, as it is not tied to particular file formats, acquisition systems, or data modalities. At its back end, DataJoint is powered by the flexible and scalable open-source relational database management system MySQL (or its equivalents such as MariaDB). However, DataJoint users do not need to learn SQL. They can manipulate data transparently through two of the most popular languages for scientific data analysis: Python and MATLAB. Data can be concurrently accessed and manipulated by multiple users using either language, or distributed across multiple computers for parallel processing.

## Using DataJoint

### Defining data elements

Consider a particular real-world neuroscience study comprising a series of experiments that involve simultaneous *in vivo* recordings of several types of physiological signals (wholecell membrane potential or V_m_, local-field potentials or LFP, and calcium fluorescence signals), stimuli (visual display and optogenetic stimulation of targeted neuronal populations), and behaviors (locomotion, whisking, eye movements) [7]. Figure 1 A illustrates such an experiment.

**Figure 1:**
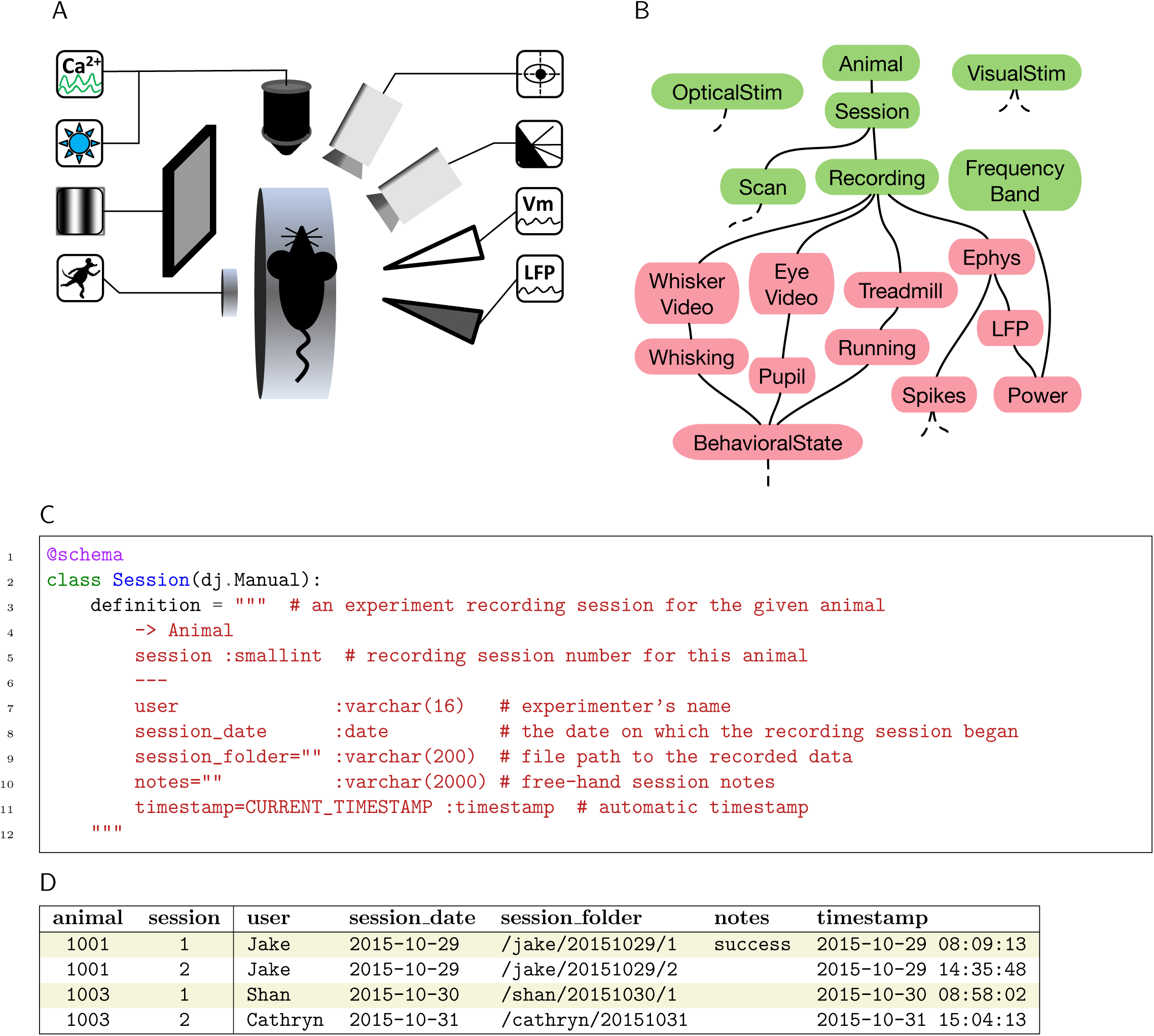
An example experiment and its DataJoint schema. A. A neuroscience experiment with multiple stimulation and acquisition modalities (counterclockwise from top left corner): fluorescence imaging of calcium signals (Ca^2+^), light stimulation of optogenetic probes, visual stimulus, treadmill motion recording, local-field potential recording (LFP), whole-cell patch membrane potential recording (V_m_), video of whisker movements, video of eye movements. **B.** The entity relationship diagram (ERD) of a DataJoint schema comprising base relations storing externally entered data (green) and automatically populated data (red). **C.** The Python class for the base relation `Session` specifying the relation’s heading. A dependency on `Animal` is indicated with the arrow `->.` An additional primary key attribute, `session,` enables multiple sessions per animal. Dependent attributes are separated from primary key attributes by ---. Each attribute has a name, an optional default value, a datatype, and an optional comment. **D.** Example contents of `Session.` The vertical divider separates the primary key attributes ‘`animal`’ and ‘`session`‘ from the dependent attributes.

To work with data from such studies, DataJoint users create a *schema* (or several schemas) comprising a collection of *base relations* to represent the various elements of the experiments (Fig. 1B). Relations are DataJoint’s basic data representation and can be thought of as simple tables with a *heading* and a *body.* The heading specifies attribute names and datatypes. The body comprises a set of *tuples* of attribute values. Base relations are stored in the database whereas *derived relations* may be constructed from base relations for data queries. For detailed definitions of the Relational Data Model, see Table 1.

**Table 1:**
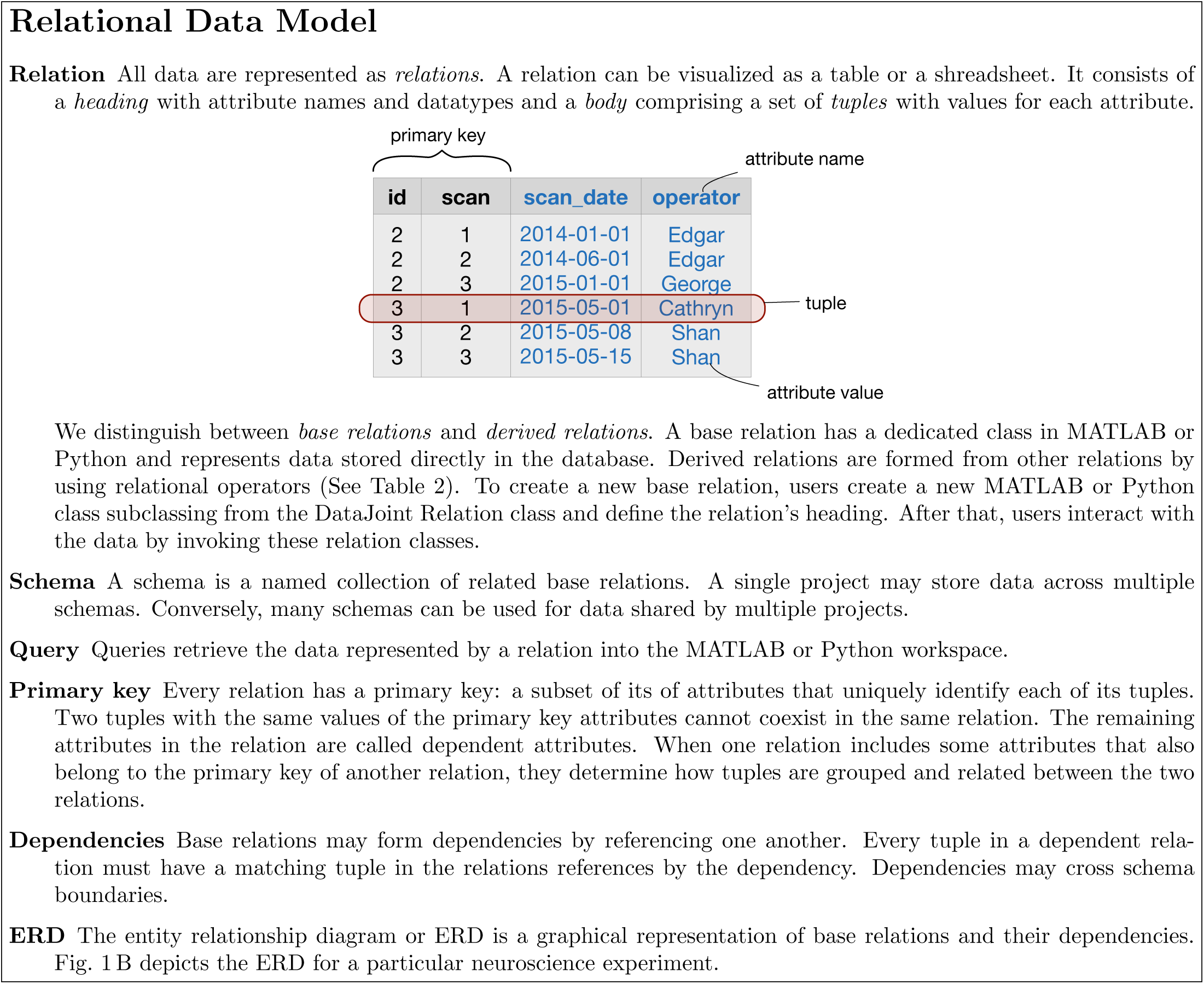
Key concepts of the relational data model as used in DataJoint.

A base relation is created in the form of a class in MATLAB or Python (Fig. 1 C) that defines the relation’s heading. The heading definition comprises a description of the relation, dependencies on other base relations, and a set of *attributes.* Each attribute has a name, an optional default value, a datatype, and an optional description. Attributes that comprise the *primary key* of the relation are separated from the remaining attributes by the divider ---.

DataJoint provides all the functionality for accessing and manipulating data through the base relation classes.

### Defining dependencies

The database must respect dependencies between data elements and prevent incomplete, orphaned, or mismatched data. DataJoint facilitates setting, enforcing, and displaying data dependencies. The edges of the graph in the entity relationship diagram (ERD) in Fig. 1 B denote dependencies directed downward: relations below are dependent on relations above when connected. Chains of dependencies effectively set the order in which data must be populated. Thus the ERD serves as an effective communication tool for the overall data organization and the sequence of steps to be followed for data entry and processing. Each base relation can depend on multiple other relations but the dependency graph must be acyclic: a relation cannot depend on itself or on other relations that depend on it directly or through other relations.

To create a dependency, the dependent relation’s data definition must include the line `-> Reference,` where `Reference` is the class name of the referenced base relation.

Setting a dependency has two effects:

1. the primary key attributes of the referenced relation are copied into the definition
2. a foreign key constraint is created to the referenced relation.

The foreign key constraint causes the database to reject any new tuple in the dependent relation unless there exists a matching tuple in the referenced relation. Conversely, deleting a tuple from the referenced relation will cause all matching tuples in all the dependent relations to be deleted too.

For example, when `Session` depends on `Animal` (Fig. 1B), `Animal’s` primary key attribute `animal` is automatically included in `Session’s` heading (Fig. 1 C and D). A new session cannot be entered for an animal that has not yet been entered; and when an animal is deleted, all its sessions will be deleted as well, along with all the dependent data below in the hierarchy.

Importantly for data dependencies, DataJoint treats tuples in relations as indivisible; dependencies are established between whole tuples rather than between attribute values. DataJoint methods modify relations only by inserting or deleting entire tuples and cannot update individual attribute values independently.

Such discipline guarantees that any changes of attribute values will trigger recomputation of all dependent data. Of course, users can deliberately intervene and modify values manually to bypass dependencies when necessary, provided that they have been granted update privileges by the database administrator.

The primary keys and dependencies between base relations allow defining a rich variety of relationships between data elements. Three common types of relationships illustrate this point (Figure 2).

**Figure 2:**
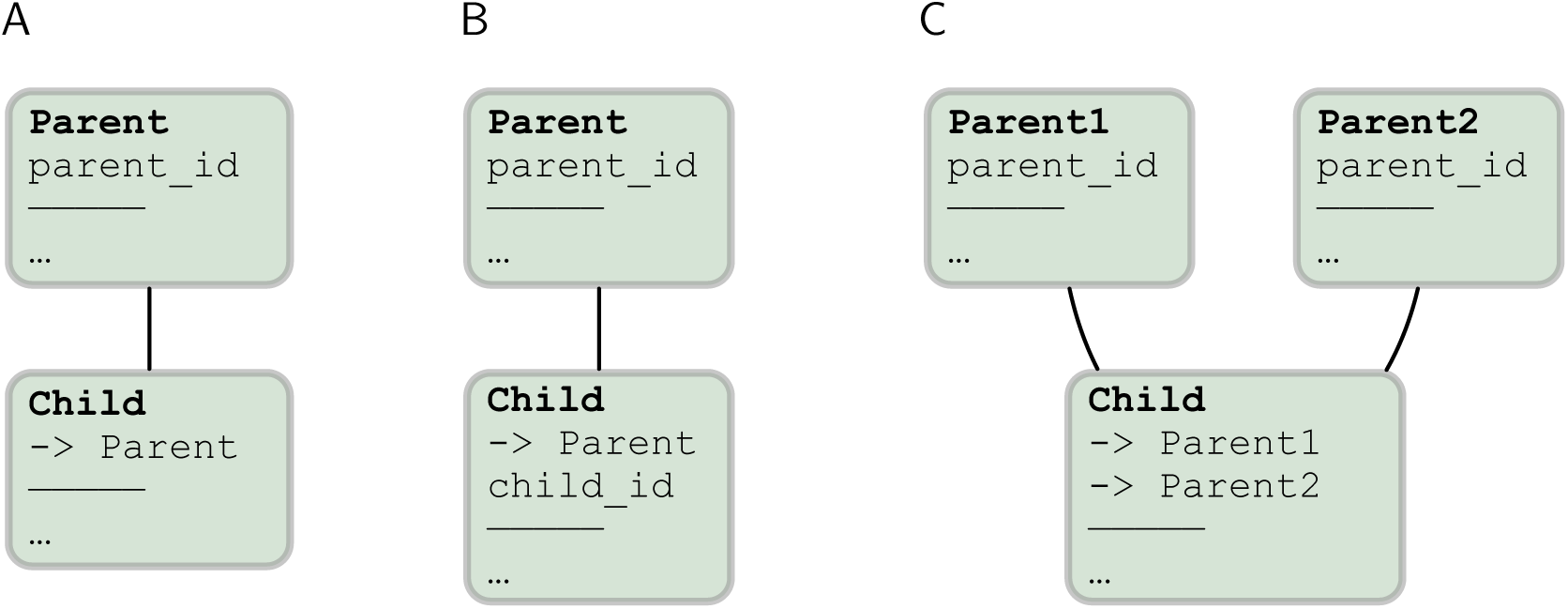
Three common replationships defined through the dependencies and primary keys of base relations. A. **A** one-to-one relationship. **B.** A one-to-many (hierarchical) relationship. **C.** A combinatorial relationship.

- In a one-to-one relationship (Fig. 2 A), relation `Child` declares a dependency within its primary key on `Parent` but does not add any new attributes to its primary key. Thus the primary key for `Child` is the same as for `Parent:` only one tuple in `Child` can exists for each tuple in `Parent`.
- In a one-to-many relationship (Fig. 2 B), relation `Child` declares a dependency within its primary on `Parent` but also declares an additional attribute `child_id` in its primary key, which allows `Child` to have multiple tuples matching each tuple in `Parent`.
- In a combinatorial relationship (Fig. 2C), relation `Child` declares a dependency within its primary key on multiple relations. In this example, `Child’s` primary key combines the primary key attributes from both `Parent1` and `Parent2.` This means that `Child` holds tuples corresponding to any combination of tuples in `Parent1` and `Parent2`.

Examples of other types of relationships may be found in the online resources.

### Querying data

DataJoint provides a minimal yet powerful set of operators on relations: restriction, projection, and join. These operators allow transforming relations into new derived relations. Table 2 summarizes these operators.

**Table 2:** Relational operators of DataJoint

| id | species | sex | date_of_birth |
| --- | --- | --- | --- |
| 1 | mouse | F | 2015-04-01 |
| 2 | monkey | M | 2011-12-01 |
| 3 | mouse | M | 2015-05-08 |
| 4 | mouse | F | 2015-05-08 |

| id | scan | image |
| --- | --- | --- |
| 2 | 1 | (BLOB) |
| 2 | 2 | (BLOB) |
| 2 | 3 | (BLOB) |
| 3 | 1 | (BLOB) |
| 3 | 2 | (BLOB) |
| 3 | 3 | (BLOB) |

| id | species | sex | date_of_birth |
| --- | --- | --- | --- |
| 2 | monkey | M | 2011-12-01 |
| 3 | mouse | F | 2015-05-08 |

| id | species | sex | date_of_birth |
| --- | --- | --- | --- |
| 1 | mouse | F | 2015-04-01 |
| 4 | mouse | F | 2015-05-08 |

| id | scan | species | sex | date_of_birth | image |
| --- | --- | --- | --- | --- | --- |
| 2 | 1 | monkey | F | 2011-12-01 | (BLOB) |
| 2 | 2 | monkey | F | 2011-12-01 | (BLOB) |
| 2 | 3 | monkey | F | 2011-12-01 | (BLOB) |
| 3 | 1 | mouse | M | 2015-05-08 | (BLOB) |
| 3 | 2 | mouse | M | 2015-05-08 | (BLOB) |
| 3 | 3 | mouse | M | 2015-05-08 | (BLOB) |

| id | scan | date_of_birth | image |
| --- | --- | --- | --- |
| 2 | 1 | 2011-12-01 | (BLOB) |
| 2 | 2 | 2011-12-01 | (BLOB) |
| 2 | 3 | 2011-12-01 | (BLOB) |
| 3 | 1 | 2015-05-08 | (BLOB) |
| 3 | 2 | 2015-05-08 | (BLOB) |
| 3 | 3 | 2015-05-08 | (BLOB) |

$\text{.pro('id' -> animal\_id', 'image')} =$
| animal_id | scan | image |
| --- | --- | --- |
| 2 | 1 | (BLOB) |
| 2 | 2 | (BLOB) |
| 2 | 3 | (BLOB) |
| 3 | 1 | (BLOB) |
| 3 | 2 | (BLOB) |
| 3 | 3 | (BLOB) |

The output of each relational operator is a proper relation in its own right with its primary key, uniquely named attributes, and the full range of data query methods. This property, called *algebraic closure,* allows expressings highly specific queries from existing queries intuitively and laconically.

The starting point of any relational expression are base relations represented by their classes. For example, after executing the following assignment in either Python or MATLAB,

~~~
rel = Ephys()
~~~

the variable `rel` will represent the contents of the base relation `Ephys,` which represents electrophysiological recordings: local field potentials and spikes in our schema (Fig. 1 B).

Restriction (represented by the logical AND operator `&)` selects a subset of tuples based on some condition. For example, `Animal` may be restricted by a structure specifying values of attributes `species` and `sex,` in MATLAB,

~~~
r.species = ‘mouse’
r.sex = ‘M’
~~~

or in Python

~~~
r = dict(species=’mouse’, sex=’M’)
~~~

to produce the relation containing all male mice,

~~~
male_mice = Animal() & r
~~~

Relations can be restricted by conditions in the form of character strings, structures or structure arrays, or other relations. For example, other relations may be restricted with `male_mice` even in combination with other restrictions:

~~~
rel = Ephys() & male_mice & ‘sampling_rate > 10000’
~~~

The result `rel` will represent all Ephys recordings from male mice with acquisition sampling rates above 10 kHz when `sampling_rate` is an attribute of `Ephys`.

As another example, the relation `Running` contains episodes of the animals’ locomotion inferred from treadmill sensor recordings in relation `Treadmill.` Then the restriction

~~~
rel = Session() & Running()
represents all sessions with at least one episode of running.
~~~

Restrictions can also take the negative form using the – (minus) operator. For example,

~~~
rel = (LFP() & Treadmill()) - Running()
~~~

contains all LFP recordings in sessions that included treadmill recordings but no running episodes where found.

The join operator (**\***) produces a relation comprising all possible combinations of matching tuples from its two argument relations.

For example, the relation

~~~
rel = Spikes() * LFP()
~~~

will contain both spiking and LFP data for the same `Ephys` recordings.

Relational operators can be combined to produce highly specific expressions. For example,

~~~
rel = Spikes() * LFP() & (Animal() * Session() & ‘datediff(session_date, date_of_birth)<=28’)
~~~

is similar to the previous example but the result is restricted to cases when the animals were 28 days old or younger at the start of the recording session. Since DataJoint passes its restriction conditions to SQL, restrictions can call SQL functions such as `datediff` here for computing the difference between the dates. Note that `session_date` comes from `Session` whereas `date_of_birth` comes from `Animal.` Thus the relation `Animal*Session` has both dates required for calculating the age.

The *projection* operator allows selecting and renaming attributes as well as computing new attributes, including summary statistics on other relations. Please refer to the online documentation for additional details.

Relation objects are only symbolic representations of the data and relational expressions are only symbolic manipulations. Once the desired relation is formulated, the actual data are retrieved from the database into a structure array using the `fetch` method

~~~
data = rel.fetch()
~~~

### Entering and computing data

DataJoint distinguishes between *manual* and *automated* base relations. In the entity relationship diagram (Fig. 1 B), manual base relations are displayed as green nodes whereas automated base relations are displayed in red.

Manual base relations contain data entered by the experimenter or by acquisition software. They store data that are derived from external sources and are typically at the head of the dependency hierarchy. Users commonly edit manual relations directly in the form of a spreadsheet using third-party interfaces such as MySQL Workbench, Navicat, SequelPro, HeidiSQL, and others.

Automated base relations are filled automatically from MATLAB or Python with the help of their `populate` method. For example, Figures 3 and 4 list the complete implementation of the `Power` base relation, which computes the average power of the LFP signal for various frequency bands in our schema (Fig. 1 B).

**Figure 3:**
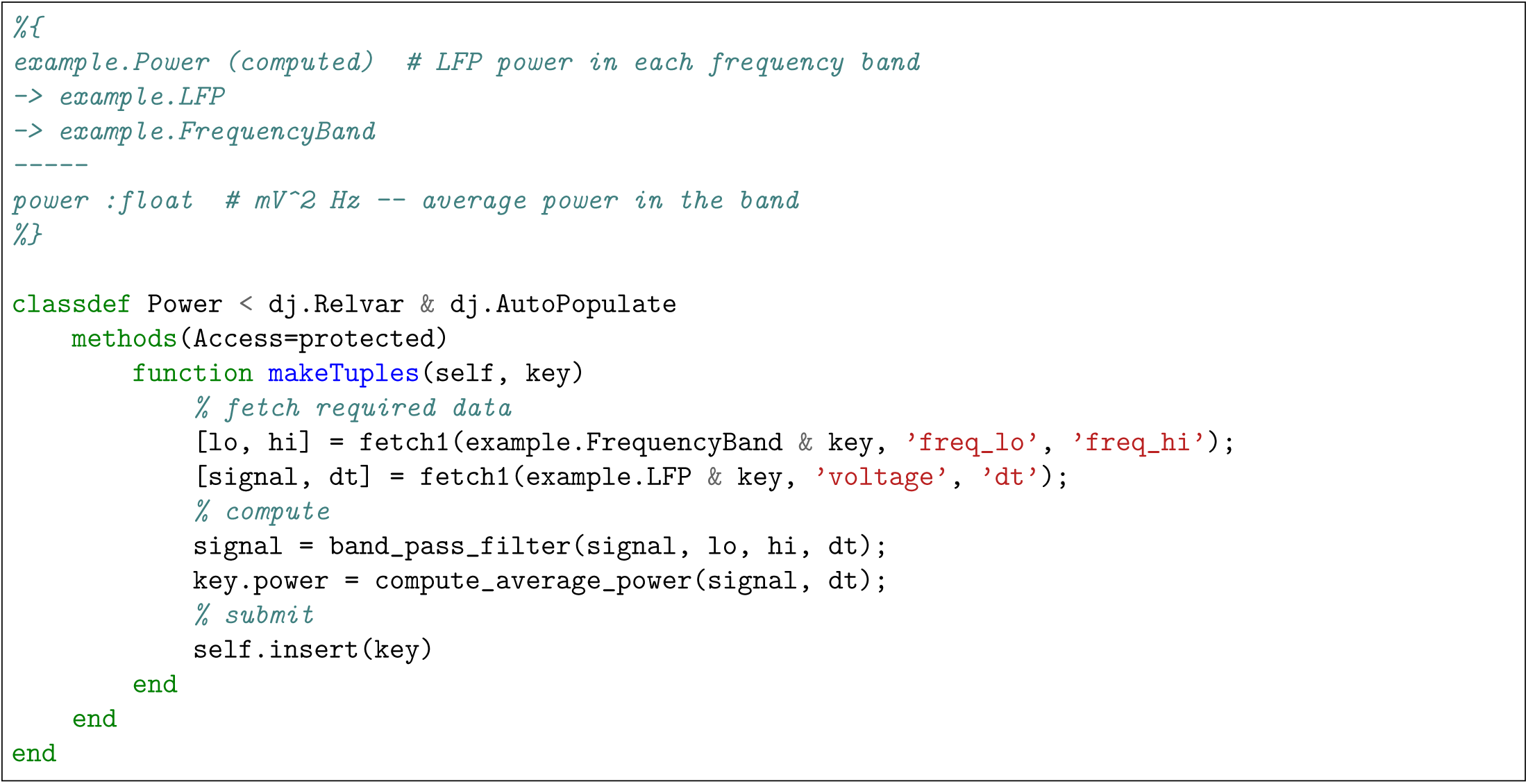
The class for the base relation `Power` in MATLAB.

**Figure 4:**
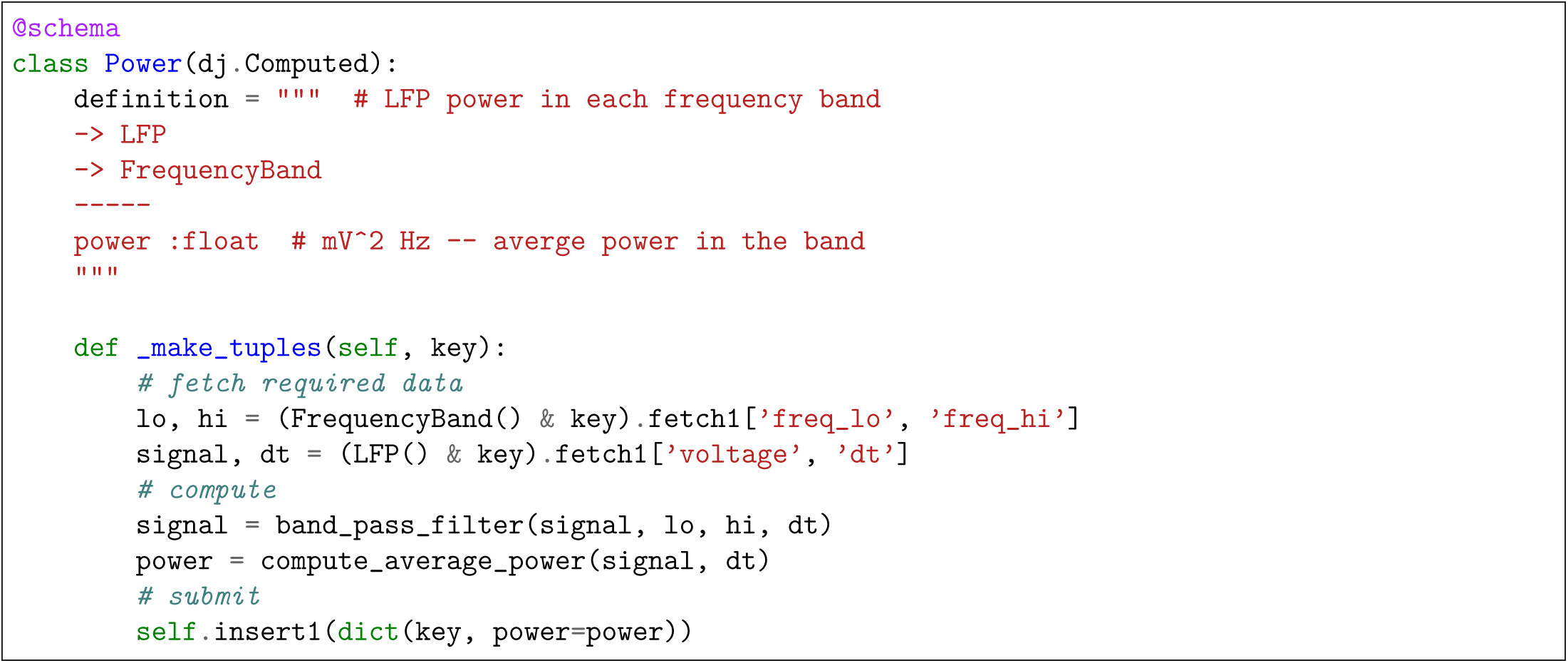
The class for the base relation `Power` in Python.

Execution of the following commands will fill `Power` for all available data.

~~~
rel = Power()
rel.populate()
~~~

The `populate` method always “knows” what needs to be computed using the base relation’s dependencies. It compares the contents of the base relation to those of its immediate neighbors upstream in the dependency hierarchy. The job list is defined as the join of the immediate upstream neighbors of the populated relation *minus* the population itself.

For example, `Power` depends on `LFP` and `FrequencyBand`. Then the restricted join

~~~
missing = LFP() * FrequencyBand() - Power()
~~~

will express all combinations of tuples in `LFP` and `FrequencyBand` for which `Power` does not yet have any entries. Each tuple in `missing` specifies an isolated job to be performed for `Power.` This logic is implemented internally and is provided here only to help users understand what happens under the hood of a `populate` call.

When `rel.populate()` is called, it executes `rel.makeTuples(key)` (in MATLAB) or `rel._make_tuples(key)` (in Python) for the primary key value of each tuple in `missing`.

Users specify the computation of new tuples for each item in `missing` using the *make-tuples* callback method (Figures 3 or 4), which consists of three parts:

1. fetch the required data from other relations upstream in the dependency hierarchy, always restricting by the argument `key`,
2. use fetched data to compute attributes of the relation that are missing in `key`,
3. create new tuples that combine the newly computed attributes and `key` and submit them to the database using the `insert` method.

Each *make-tuples* call runs inside an isolated *transaction* so that its results do not become visible to other processes until the entire call completes successfully. If an error occurs during a *make-tuples* call, any partially populated data are discarded and never become visible to downstream computations.

The `populate` method has several options to control its behavior. In particular, it has the option of using the built-in *job reservation* process to enable efficient distributed execution. With job reservation enabled, users simply execute `populate` on multiple computers to run the *make-tuples* jobs in parallel without conflicts. Please refer to the online documentation for `populate` options and various techniques for customizing the processing chains.

### Sharing data and distributed computation

Through the features described above, DataJoint naturally supports collaboration and distributed access by

- setting, enforcing, and communicating dependencies between data elements,
- keeping data definition and computation code together in the same class for easy review and sharing,
- storing and organizing data in a popular database management system (MySQL) that can be accessed through a variety of third-party interfaces,
- allowing safe simultaneous access to the same data through standard transaction processing provided by MySQL, and
- providing an automatic and transparent job reservation process for conflict-free distributed processing.

## Discussion

The relational data model has served as the theoretical foundation for the majority of mainstream database systems for over forty years and has been shown to be the most cogent and principled approach for representing and manipulating data of arbitrary complexity [10]. Yet, relational organization of data in science labs is still uncommon. This disconnect is partly due to SQL’s position as the only wide-spread implementation of the relational model. SQL and its dialects have deviated substantially from the simplicity of the relational data model and have overgrown with extraneous complexity.

Object-relational mappers (ORM) are software tools that map objects in computer memory to persistent storage such as relational databases. Since DataJoint constructs objects with persistently stored data, it can be classified as an ORM. Several other object-relational mappers are available for Python: SQLAlchemy, Django ORM, Peewee, PonyORM, SQLObject. They take a similar approach of converting objects and idioms of the host programming language into SQL queries for processing by the database server. We are not aware of any ORM tools for MATLAB besides DataJoint.

DataJoint addresses other needs than ORMs: it is specifically designed for providing a robust and intuitive data model for scientific data processing chains. As such, it does not attempt to simply mirror the features and capabilities of SQL. Instead, DataJoint imposes constraints and conventions to achieve the expressive power and simplicity of queries by strict adherence to the relational data model. In science, both the structure of the data and the queries evolve frequently. A simple, sound data model is of greater importance than in other database applications.

Some of the distinct constraints imposed by DataJoint include the following:

1. All data are represented as proper relations with a primary key and uniquely named attributes. This applies to base and derived relations. Relational operators follow consistent rules for determining the primary key of its result. As a result, DataJoint’s operators are algebraically closed, allowing building complex expressions from simpler expressions.
2. Data in base relations are updated only by inserting or deleting entire tuples: updates of attribute values are not supported. As discussed in the text, this limitation is necessary because referential constraints (foreign keys) enforce data dependencies only between tuples and not between individual attribute values.
3. Dependencies between base relations are acyclic, *i.e.* they cannot form loops. This restriction simplifies data definition but, perhaps counter-intuitively, does not prevent specifying arbitrary relationships between data elements, including directed graphs with cyclic dependencies, for example.
4. DataJoint limits relational operators to enforce clarity. For example, the projection operator (see Table 2 and online documentation) does not allow projecting out the primary key attributes. Consequently, the resulting relation has the same number of tuples as the original relation and every tuple is unique. If the user does intend to derive a relation with a different primary key, she must explicitly declare a base relation with this primary key and use it to formulate the proper query. In practice, this is not a real limitation but a specific prescription of how data must be defined and manipulated in a uniform and explicit manner.
5. Foreign keys always link identically named attributes in both relations. This convention simplifies the specification of dependencies and of relational operators. For example, a single join operator in DataJoint can perform the same work as the multiple forms and parameterizations of the JOIN operators in SQL. This convention is particularly important in DataJoint because it allows direct logical linking of relations separated by many intermediate dependencies. In a large schema, this convention may lead to long composite primary keys low in the dependency hierarchy, but these are efficiently handled by MySQL’s storage engines.

DataJoint’s restricted relational data model represents a conceptual shift in database interactions: In SQL queries, users explicitly enumerate and match individual attributes. In contrast, DataJoint users formulate dependencies and queries at the level of entire relations. As a result, DataJoint’s fast, intuitive, and expressive data definition and manipulation languages enable scientists to flexibly adapt their data processing chains to evolving demands.

### Examples and resources

To benefit from DataJoint, users need not appreciate the various technical considerations underlying its capabilities. We encourage interested readers to review the online documentation to get up and running with DataJoint quickly. For tutorials, a gallery of working schemas for MATLAB and Python, documentation, and references to scientific papers using DataJoint, please visit http://datajoint.github.com.

